# Sustainable and Ecofriendly Nanoparticle Synthesis Using Microbial Strains Isolated From Heavy Metal Rich Industrial Effluent

**DOI:** 10.1101/2024.03.16.585342

**Authors:** Soshina Nathan, Soumya Mathunny, J Anjana

**Affiliations:** Fatima Mata National College (Autonomous), Kollam, Kerala, India

**Author notes:** Corresponding Author: Dr. Soshina Nathan, PG and Research Department of Zoology, Fatima Mata National College (Autonomous), Kollam, Kerala, India.

**Keywords:** heavy metal, monodisperse, nanoparticles, sustainable, microbes, Titanium industry effluent

## Abstract

Given the enormous potential of metal nanomaterials, their sustainable production is of paramount importance and is a key area of focus worldwide. In this regard, bacteria are highly valued because of their potential for rapid, cost-effective and eco-friendly metal nanomaterial synthesis. In this study, culture supernatants of *Bacillus cereus* and *Curvularia* sp isolated from heavy metal rich Titanium industry effluent effectively synthesised cobalt and copper nanoparticles of narrow size range at room temperature, neutral pH and static conditions within 2-7 days. This was verified by visible colour changes, UV-Vis spectroscopy and FT-IR. The UV-Visible spectra of the biosynthesized cobalt and copper nanoparticles exhibited sharp narrow peaks at 341 and 342 nm. This suggested that the cobalt and copper nanoparticles were not only small but also had a narrow size distribution, a feature rarely reported in biosynthesis studies. Furthermore, our approach was conducted at room temperature using cell-free supernatant, eliminating the need for additional heating or cooling, and minimising processing thus making the process energy-efficient, cost effective and sustainable. This is a first report on the production of monodisperse cobalt and copper nanoparticles by microbes isolated from this novel extreme environment.

**Graphical abstract:** 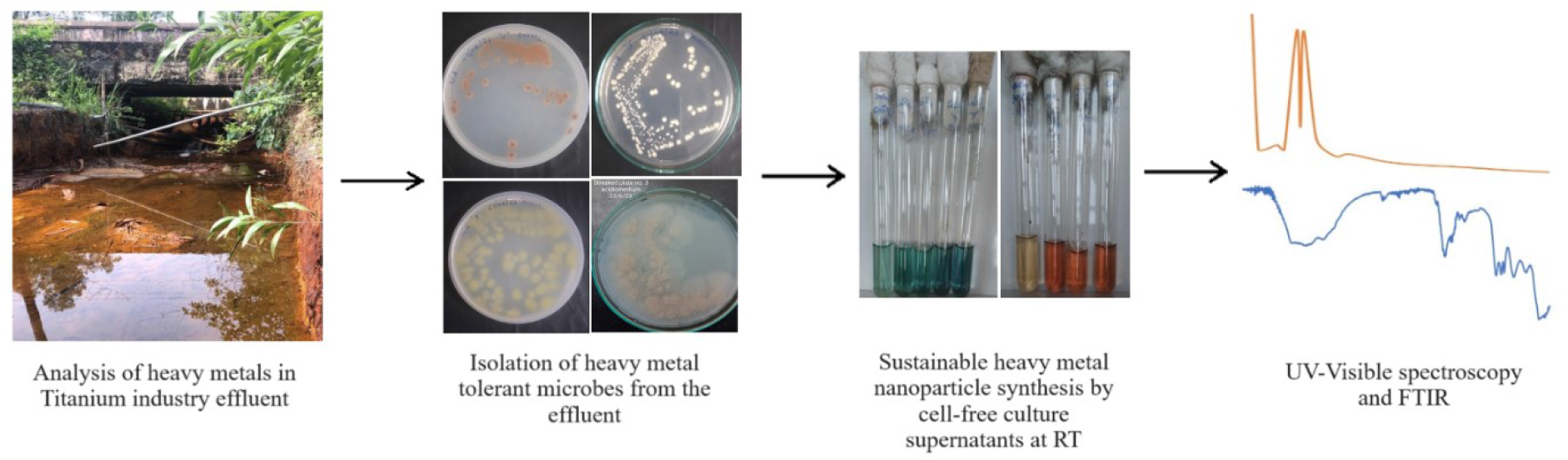

## 1. INTRODUCTION

In the rapidly evolving field of nanotechnology, the synthesis of nanoparticles has become a subject of significant interest due to their unique properties and wide range of applications. However, conventional methods of nanoparticle synthesis often involve the use of toxic chemicals and high energy consumption, posing serious environmental concerns. Microorganisms have long been known for their ability to interact with metals in their environment, with some species demonstrating remarkable resistance to heavy metal toxicity. These resistance mechanisms include permeability barriers, metal sequestration, where the organism stores the metal in a form that is less toxic; metal reduction, where the organism changes the oxidation state of the metal to a less toxic form; and metal efflux, where the organism actively pumps the metal out of its cells. Some microorganisms can even utilize heavy metals for their metabolic processes, effectively ‘cleaning up’ the environment through a process known as bioremediation [1]. These resistance mechanisms can be harnessed for the synthesis of nanoparticles. By isolating microbial strains from industrial effluent rich in heavy metals, we can tap into a naturally occurring, eco-friendly method of nanoparticle synthesis.

Titanium extracted by the chloride process produces effluent that is rich in heavy metals including cobalt, copper, chromium etc [2]. The application of heavy metal-resistant or tolerant bacteria in the biosynthesis of metal nanoparticles is an emerging field of research with significant potential as chemical and physical methods of synthesis are costly and non-ecofriendly. There is a growing emphasis on developing an eco-friendly and green approach for producing metal nanoparticles using biological systems. While plants, fungi and algae have been used for nanoparticle synthesis, bacteria offer benefits such as easy culturing, short generation times and excellent nanoparticle stability [3]. The production of nanoparticles by bacteria can be done by three methods: by intact cells, by cell-free supernatants or using bacterial cell extracts [4]. However, the use of cell-free supernatants for nanoparticle synthesis is attractive as it simplifies downstream processing, offers scalability and eliminates the need for intact cells.

Nanoparticles of cobalt and copper are attracting increased interest because of their diverse applications in agriculture, biotechnology, chemistry, and medicine. For example, cobalt oxide nanoparticles find applications in catalysis, energy storage, as antimicrobial agents, as drug delivery systems and as contrast agents [4, 5] while copper nanoparticles are used in catalysis, electronics, sensors and have antimicrobial and anticancer activities [6]. Monodisperse nanoparticles, due to their uniform size and shape, find extensive use in various fields such as drug delivery, electronics, catalysis, and optics. Although numerous synthesis methods for monodisperse particles have been proposed, they often come with challenges. These include issues with controlling particle size, overuse of solvents and surfactants, and a lack of adaptability to changes in experimental conditions. Consequently, there is a pressing demand for a cost-effective and ecofriendly method for synthesizing monodisperse nanoparticles to meet existing and future market requirements [7].

The focus of this article is the exploration of a novel, eco-friendly approach to nanoparticle synthesis using microbial strains isolated from heavy metal-rich industrial effluent. This method leverages the inherent capabilities of these microbial strains to reduce metal ions and form nanoparticles, offering a green and sustainable alternative to traditional chemical synthesis methods. In the present study, bacteria and fungi were isolated from heavy metal-rich Titanium industry effluent as a novel sampling point. After screening, the microbes were investigated for their ability to synthesise nanoparticles of cobalt and copper extracellularly at room temperature and neutral pH using only the culture supernatants in a simple, inexpensive and ecofriendly approach. The nanoparticles of cobalt and copper were verified by UV-Visible spectroscopy and Fourier Transform Infra Red Spectroscopy and found to be monodisperse.

## 2. MATERIALS AND METHODS

### 2.1. Sample collection and processing

Untreated effluent, treated discharged effluent and sediment samples from different locations in and around a Titanium mineral processing factory in Kollam district, Kerala were collected in sterile bottles and processed for heavy metal analysis and microbial isolation within 3 hours. The physical characteristics of the different effluent samples, such as colour, pH and temperature were noted on site. The concentrations of cobalt, copper, iron, manganese and zinc in the treated discharged effluent were determined by standard methods as shown in Table 1. These metals were specifically chosen for analysis as their nanoparticles are known to have various applications.

**Table 1:** Concentration of heavy metals in the discharged Titanium industry effluent and comparison with permissible limits of these heavy metals in drinking water.

| S. No. | Heavy metal | Test method | Concentration in effluent (mg/L) | BIS (2012)- (IS 10500: 2012- second revision) (mg/L) | WHO (2022) guideline value (mg/L) | USEPA (2018) (secondary drinking water regulations) (mg/L) |
| --- | --- | --- | --- | --- | --- | --- |
| 1 | Cobalt | GCKL/QS/C/7.2 /115 | 1.017 | Not mentioned | 0.05* | Not mentioned |
| 2 | Copper | IS 3025 (Part 42) | 3.5 | 0.05 | 2 | 1.3 |
| 3 | Iron | IS 3025 (part 53): 2003 (UV Method) | 4.0 | 0.3 | Not of health concern at levels causing acceptability problems in drinking-water | 0.3 |
| 4 | Manganese | IS 3025-1999 | 40.0 | 0.1 | 0.08 | 0.05 |
| 5 | Zinc | IS 3025 (Part 49) | 12.2 | 5 | Not of health concern at levels found in drinking-water. | 5 |

\* Based on WHO, 2003 guidelines

### 2.2. Isolation of microbes

To isolate microbes, 10 ml or 10 g of the well mixed effluent or sediment sample was vortexed with 90 ml of autoclaved distilled water to make a 10^−1^ dilution. Serial dilutions of the samples up to 10^−4^ were prepared in autoclaved distilled water and 0.1 ml of each dilution was plated on half strength nutrient agar as this was found to be more suitable for the growth of the organisms. Half strength nutrient agar was prepared by dissolving 14 g of nutrient agar powder (Sisco Research Laboratories), instead of 28 g, in 1 litre of water and autoclaved. The inoculated plates were then incubated at room temperature for 24 - 48 hrs. Eight morphologically different microbial colonies were randomly selected and then sub-cultured and re-streaked several times to purify the cultures. The pure cultures were then stored on half strength nutrient agar plates at 4 °C or as 20 % glycerol stocks at -20°C.

### 2.3. Nanoparticle synthesis and analyses

The eight isolates were then tested for their ability to form nanoparticles of the tested heavy metals that were found in the effluent. For this, 10 mM salt solutions of Ferric Chloride (FeCl_3_), Zinc sulphate (ZnSO_4_.7H_2_O), Manganese Sulphate (MnSO_4_.H_2_O), Copper Sulphate (CuSO_4_.5H_2_0), and Cobalt Chloride (CoCl_2_.6H_2_O) each were prepared and autoclaved. These were used as precursor solutions for nanoparticle synthesis. The microbial cultures were inoculated in half strength nutrient broth (Sisco Research Laboratories), and incubated at room temperature for 48 hours. The cultures were centrifuged at 4000 rpm for 20 minutes. Then, different volumes of the cell-free supernatants were mixed separately with various volumes of the 10mM salt solutions. Uninoculated half-strength nutrient broth mixed separately with 10mM salt solutions were used as controls. The mixtures of culture supernatants/uninoculated media and salt solutions were kept in the dark at room temperature under static conditions for up to seven days for nanoparticle synthesis. During this period, any colour change observed in the mixtures indicated that the microbes were nanoparticle producers. Conversely, if the colour of the salt solution remained unchanged following inoculation with culture supernatants, the microbes were considered non-producers of nanoparticles. UV-Vis spectroscopy, being one of the most important and simplest methods, was employed to validate the synthesis of nanoparticles. The samples that exhibited colour changes were then subjected to UV-visible spectroscopy in a 1 cm path quartz cuvette and scanned from 200-700 nm range using a V-730 UV-Visible spectrophotometer. Dilution of the samples was done to get absorbance values in the range of 0.8-1.5 OD.

For Fourier Transform-Infra Red analyses, the mixtures were centrifuged, and the resulting particles were washed with distilled water three times to remove any water soluble metabolites or biological molecules. The particles were then washed in ethanol, dried in a hot air oven and then ground with KBr pellets for FTIR analysis. The scanning was done from 500–4000 cm^−1^ at a resolution of 1 cm^−1^ using an FTIR-Jasco V-650 Spectrophotometer.

### 2.4. Microbial Identification

Gram staining was performed on the isolates. The strains which showed the highest potential and the shortest time for nanoparticle synthesis, were identified by Matrix-Assisted Laser Desorption/Ionization Time-of-Flight Mass Spectrometry (MALDI-ToF MS) which has emerged as a rapid, reliable, sensitive and cost-effective technique for the identification of microbes [8, 9].

## 3. RESULTS

### 3.1. Physicochemical properties of Titanium industry effluent

The untreated effluent had a fluorescent green appearance, a temperature of 28°C and was extremely acidic with a pH of 0. The lime-treated discharged effluent was pale yellow with a pH and temperature of 3.1 and 26°C. Samples collected less than 200 m away from the factory also had a pale yellow colour and showed an increase in pH up to 3.5 and temperature up to 32°C.

As an initial step in our study, we measured the concentrations of heavy metals present in the collected effluent samples [Table 1]. As these metals can leach into ground water and contaminate water resources, we compared the levels of heavy metals in these collected samples with the drinking water guidelines set by three authentic standards, the Bureau of Indian Standards (IS10500: 2012), US EPA (2018) and the World Health Organization (WHO, 2022) [10, 11, 12]. The levels of Cu, Fe, Mn and Zn surpassed the acceptable levels set by the standards, with the exception of Cobalt as Cobalt isn’t included in any of these drinking water standards. However, according to the WHO guidelines from 2003 [13], the Cobalt levels were found to be 20 times higher than the permissible limit.

### 3.2. Microbiological analysis

Bacteria and fungi were isolated from the discharged effluent by spread plate method and purified by repeated streaking on half strength nutrient agar. To avoid redundancy, eight morphologically different cultures, two fungi and six Gram-positive bacteria were selected for further studies [Figure 1]. Two of the microbial isolates which showed good potential for extracellular synthesis of heavy metal nanoparticles were identified by MALDI-TOF analysis as *Bacillus cereus* and *Curvularia sp*.

**Figure 1:**
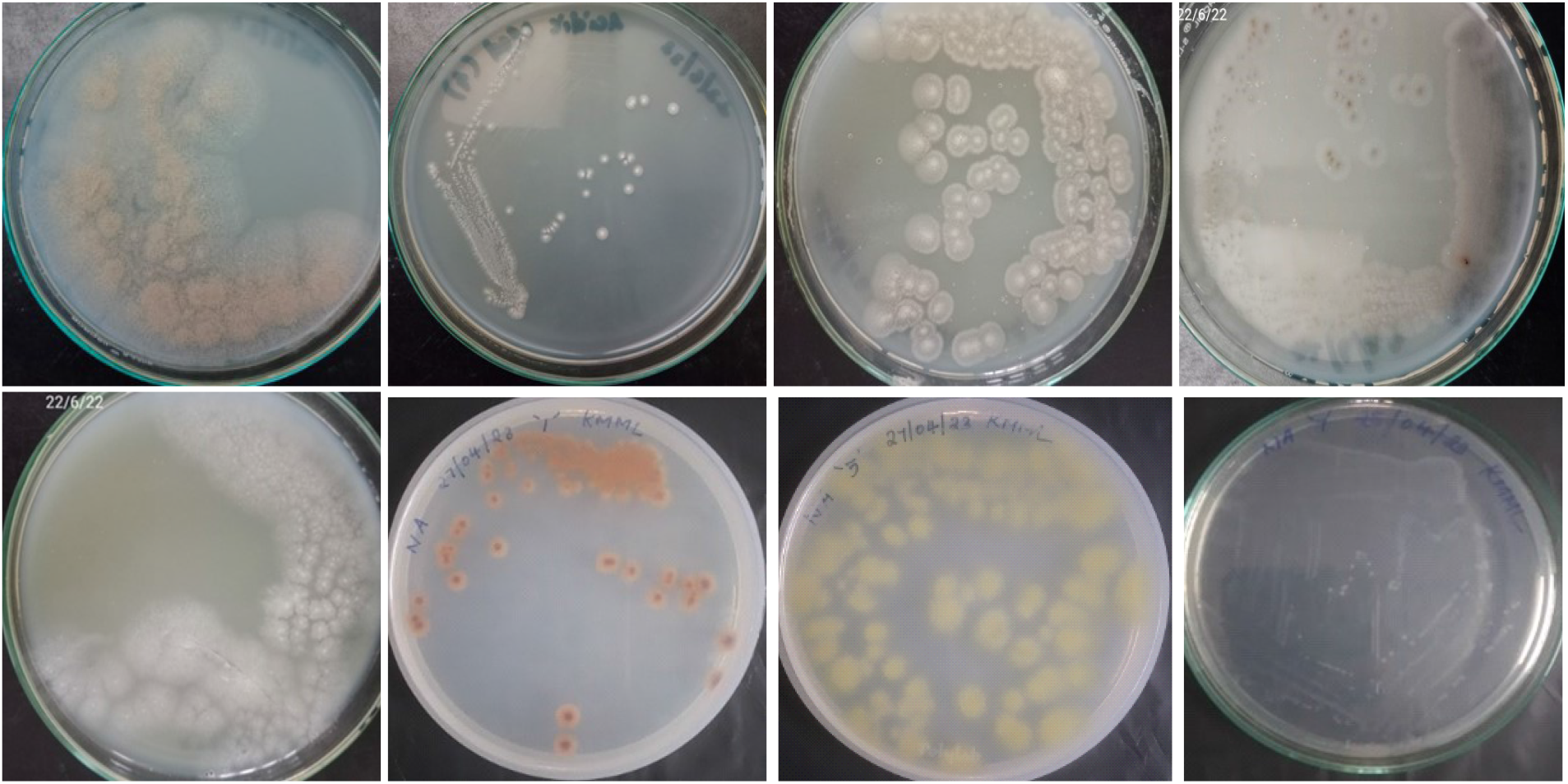
Purified microbial isolates from Titanium industry effluent on half strength nutrient agar medium.

### 3.3. Green synthesis of nanoparticles and analyses

As the microbes were isolated from heavy metal rich effluent, we reasoned that they may be able to synthesise nanoparticles of those metals. To develop a sustainable and eco-friendly approach to nanoparticle synthesis, the cell free culture supernatants of each of the eight isolates were then tested independently for extracellular production of nanoparticles of Co, Cu, Fe, Mn and Zn under static conditions at room temperature and neutral pH. The cell free culture supernatants when mixed with 10mM salt solutions of the heavy metals in a 4:1 ratio gave the best results in terms of colour change. Six out of the eight culture supernatants produced colour changes with at least one heavy metal solution. Notably, when *Bacillus cereus* culture supernatant was mixed with CuSO_4_.5H_2_O solution, the mixture changed from blue to a dark bluish green colour within 48 hrs [Figure 2a], while the *Curvularia* species culture supernatant produced a colour change from a pale brown to dark brown colour with CoCl_2_.6H_2_O within 7 days [Figure 2b].

**Figure 2:**
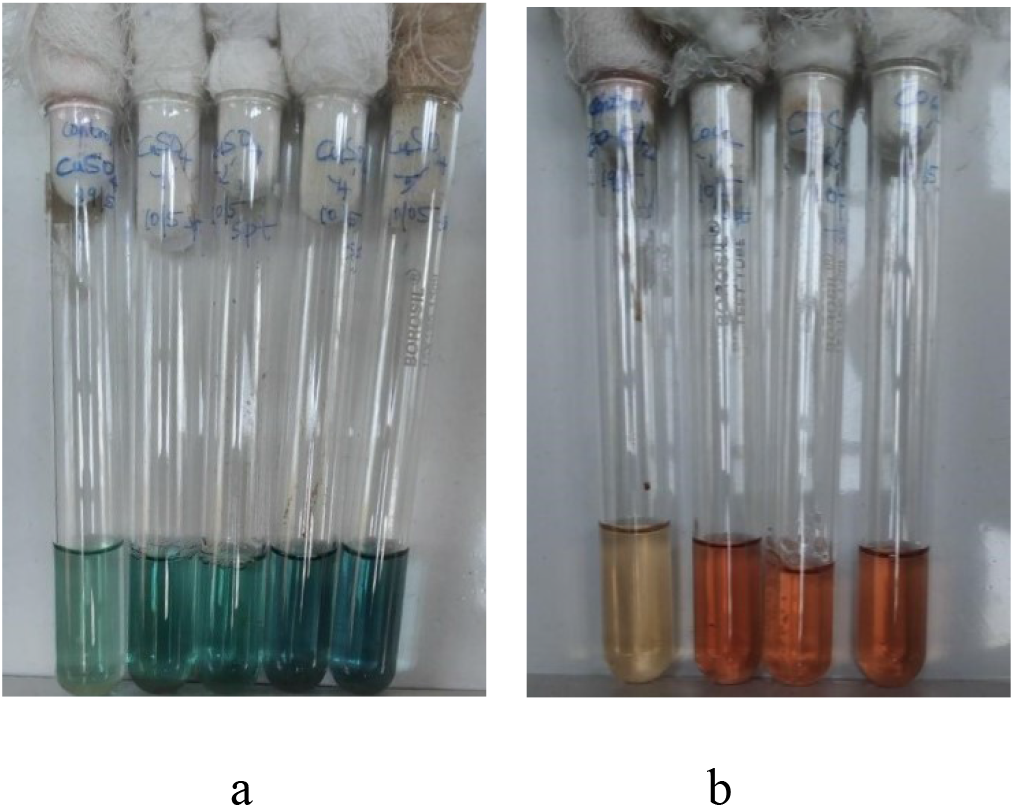
Biosynthesised Copper (a) and Cobalt (b) nanoparticles. From left to right in (a) Tube 1: Control-2mM CuSO_4_.5H_2_O solution+ Uninoculated half strength nutrient broth. Tube 2: 2mM CuSO_4_.5H_2_O solution+ Culture supernatant of isolate 1. Tube 3: 2mM CuSO_4_.5H_2_O solution+ Culture supernatant of isolate 2. Tube 4: 2mM CuSO_4_.5H_2_O solution+ Culture supernatant of isolate 4 identified as *Bacillus cereus*. Tube 5: 2mM CuSO_4_.5H_2_O solution+ Culture supernatant of isolate 5 identified as *Curvularia* species. From left to right in (b) Tube 1: Control-2mM CoCl_2_. 6H_2_O solution + Uninoculated half strength nutrient broth. Tube 2: 2mM CoCl_2_. 6H_2_O solution+ Culture supernatant of isolate 1. Tube 3: 2mM CoCl_2_. 6H_2_O solution+ Culture supernatant of isolate 2. Tube 4: 2mM CoCl_2_. 6H_2_O solution+ Culture supernatant of isolate 5 identified as *Curvularia* species.

These solutions were then subjected to UV-Vis spectroscopy in wavelengths of between 200-700 nm to determine the optical properties of the synthesized nanoparticles. The maximum absorption peaks for Copper and Cobalt nanoparticles were in the range of 341-342 nm [14, 15, 16, 17] [Figure 3 a and b]. Interestingly, the cobalt nanoparticles showed two adjacent narrow peaks at 341 and 342 nm suggesting that two sizes or types of nanoparticles may have formed.

**Figure 3:**
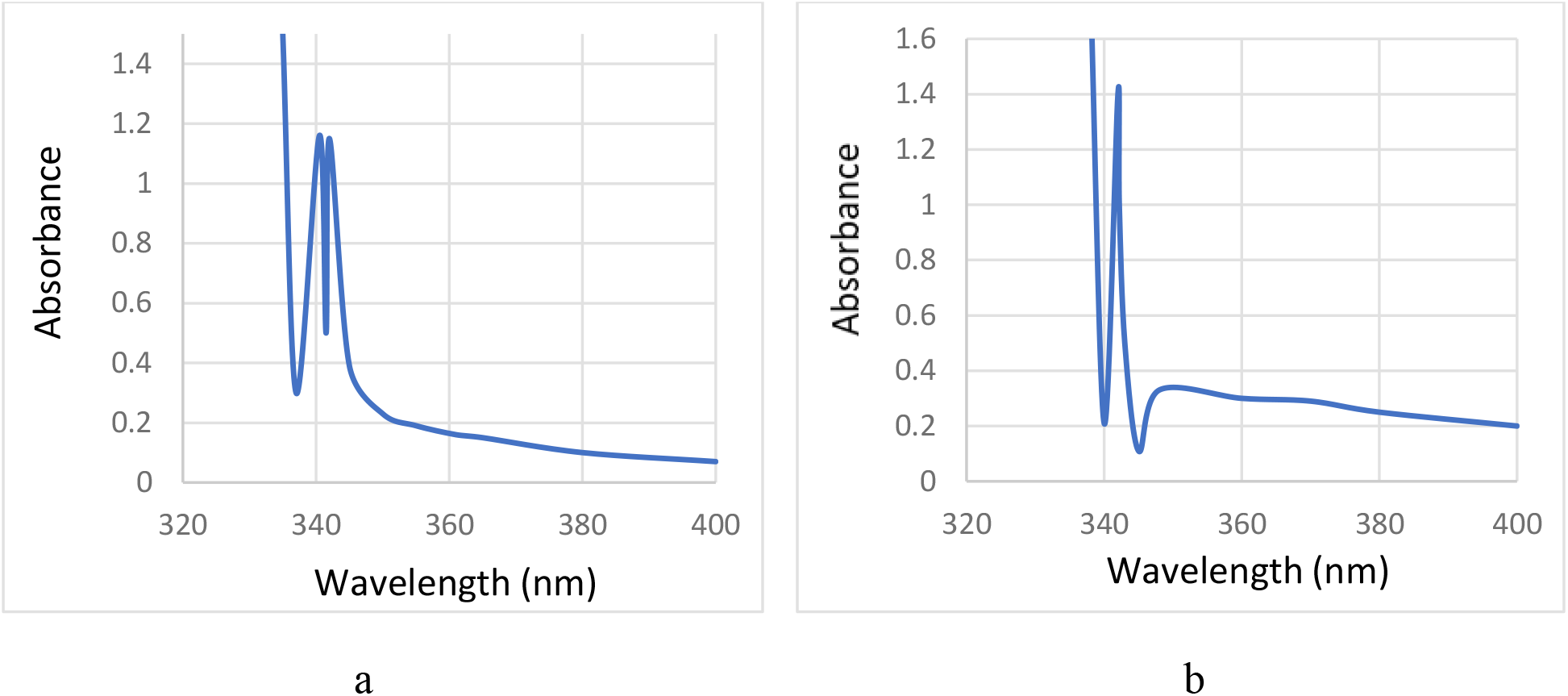
UV-Vis spectra of Co (a) and Cu nanoparticles (b) obtained from culture supernatants of the *Curvularia sp* and *Bacillus cereus* isolated from Titanium industry effluent.

Notably, over one gram of cobalt nanoparticles was produced from 25 ml of *Curvularia* culture supernatant at room temperature. To determine the possible biomolecules responsible for the reduction of the cobalt ions to nanoparticles, an FTIR analysis was done [Figure 4]. The broad peak at 3253 cm ^-1^ represented O-H stretching of carboxylic acid. The peak at 1718 cm^-1^ revealed C-O stretching which could arise from the bacteria. The presence of two peaks at 669 cm^-1^ and 528 cm^-1^ denoted Co-O stretching vibrations. The presence of carboxyl group –COOH was responsible for the capping and stabilization of the cobalt nanoparticles. These bonding features indicated the presence of Cobalt oxide nanoparticles [18, 19].

**Figure 4:**
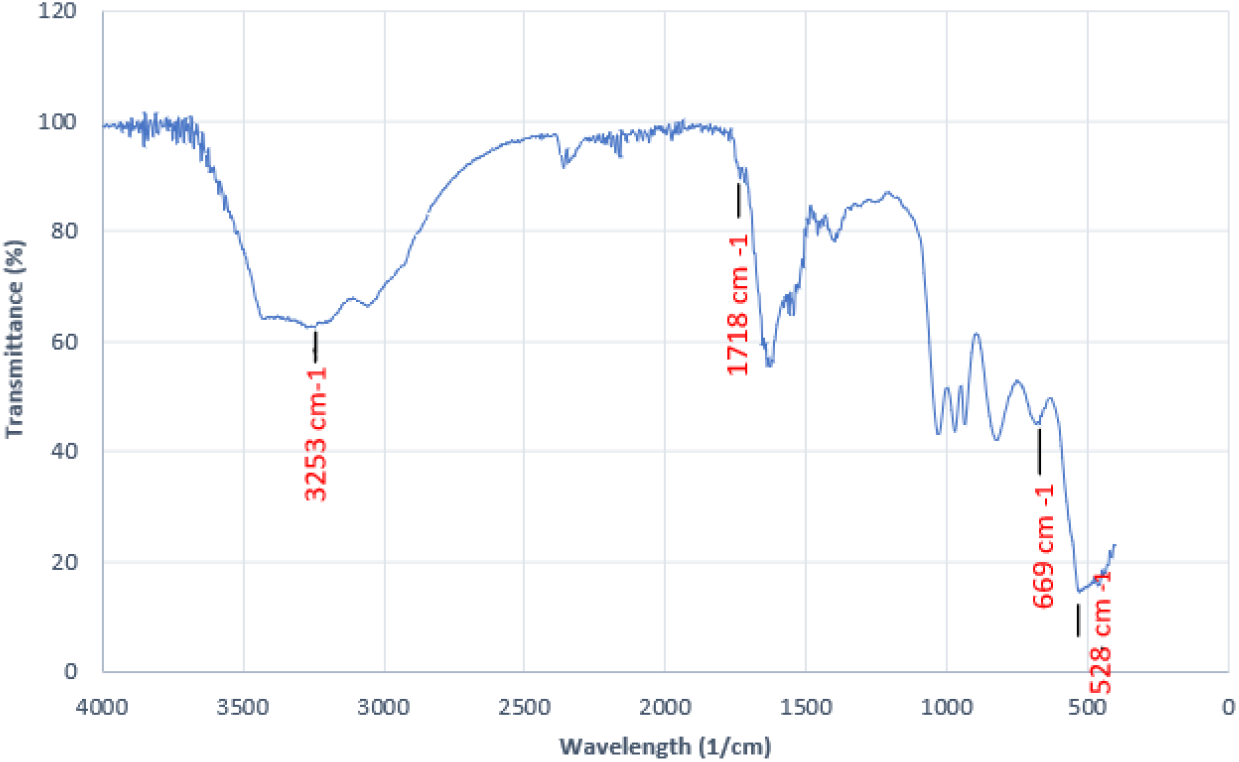
FTIR spectrum of cobalt oxide nanoparticles

## 4. DISCUSSION

To the best of our knowledge, there are no published studies that have delved into the microbial aspects of heavy metal rich Titanium industry effluents and employed the microbes present in these effluents for metal nanoparticle synthesis. Eight heavy metal tolerant microbes present in this effluent were isolated and employed for a simple and sustainable method of nanoparticle synthesis. As a lot of importance is being given to the green synthesis of nanoparticles, we tested these microbes for nanoparticle synthesis at room temperature, neutral pH and static conditions using minimal processing methods for an ecofriendly, energy and cost-effective synthesis. Two of the eight isolates were especially noteworthy in nanoparticle synthesis and were identified as *Bacillus cereus* and *Curvularia* sp by MALDI-ToF. Although, the time for production varied from 2 to 7 days depending on the culture and metal used, the use of cell free supernatants and room temperature for nanoparticle synthesis allowed for simpler processing, visualization, downstream purification and characterization of the nanoparticles.

UV-Visible spectroscopy is a simple and reliable method for monitoring the presence and stability of nanoparticle solutions [20, 21]. The peaks obtained in the UV range, about 340nm, for both the copper and cobalt nanoparticles suggested that the nanoparticles formed were of small size. It is known that smaller nanoparticles absorb light at shorter wavelengths [21]. Further, the sharp, narrow peaks of absorbance suggested that the synthesised nanoparticles were all of similar size, or monodisperse, as they absorbed a very small range of wavelengths [20]. Uniform nanoparticle size is critical, as it can significantly influence their physicochemical and biological properties. Monodisperse nanoparticles offer predictable performance, enhanced stability, and increased efficiency in applications like drug delivery and electronics [22]. It is notable that there are no published reports of monodisperse cobalt nanoparticles synthesised by bacteria. Although [23] succeeded in creating monodisperse copper nanoparticles using psychrophilic bacteria, the process necessitated synthesis at lower temperatures. When nanoparticles lose their stability, their initial extinction peak’s intensity decreases due to a decrease in the number of stable nanoparticles. Frequently, the peak will broaden or a new peak will appear at longer wavelengths [20]. In this study, the extinction peaks were stable and did not decrease in intensity or broaden even after a week, signifying that the cobalt and copper nanoparticles synthesised in this study were stable and did not aggregate. Thus, our study showcases a green and sustainable method for the synthesis of stable, monodisperse nanoparticles of cobalt and copper.

## 5. CONCLUSIONS

The culture supernatants of *Bacillus cereus* and *Curvularia* sp. isolated from heavy metal rich titanium industry effluent were used to effectively synthesise cobalt and copper nanoparticles of a narrow size range at ambient temperature, neutral pH and stationary conditions using only the water-based heavy metal salt solution and no additional reagents. This method represents a significant advancement in the field, offering a more sustainable and environmentally friendly approach to nanoparticle synthesis. This research also highlights the untapped potential of industrial effluents as sources of microbial strains capable of nanoparticle biosynthesis. Through this research, we hope to contribute to the ongoing efforts to develop green technologies and promote sustainable practices in the field of nanomaterial synthesis.

## 6. ACKNOWLEDGMENTS

The authors acknowledge the seed money sanctioned to SN for the project entitled “Isolation of Bacteria from Titanium Mineral Processing Effluent and Evaluation for Nanoparticle Synthesis” (Sanction order FC/E3/SM20/2022) provided by Fatima Mata National College (Autonomous), Kollam for carrying out this work.

## 7. AUTHORS’ CONTRIBUTIONS

SN conceived and designed the study, analysed and interpreted the data and wrote the manuscript. SM and AJ acquired and analysed the data.

## 8. FUNDING

The authors acknowledge the financial assistance provided by Fatima Mata National College (Autonomous), Kollam.

## 9. CONFLICTS OF INTEREST

The authors declare that there are no conflicts of interest.

## 10. ETHICAL APPROVALS

This study does not involve experiments on animals or human subjects.

## 11. DATA AVAILABILITY

The data used to support the findings of this study are included within the article.

## Notes

### Competing Interest Statement

The authors have declared no competing interest.

### Summary of Updates

The written content has been improved to focus on the method used for the synthesis of the nanoparticles. Also the focus is on a new aspect of the results that was not observed before- the monodispersity of the nanoparticles obtained. Further some new, more clear figures are added in this new version.

